# The PIDDosome controls cardiomyocyte polyploidization during postnatal heart development

**DOI:** 10.1101/2024.08.27.609375

**Authors:** M Leone, N Kinz, F Eichin, D Obwegs, VC Sladky, VZ Braun, D Rizzotto, L Englmaier, C Manzl, K Moos, Julia Mergner, P Giansanti, MN Garcia, MM Marques, ED Jacotot, C Savko, M Boerries, MA Sussman, A Villunger

## Abstract

The adult mammalian heart is characterized by post-mitotic polyploid cardiomyocytes (CMs). Understanding how CMs regulate cell cycle exit and ploidy can help developing new heart regenerative therapies. Here, we uncover that the PIDDosome, a multi-protein complex activating the endopeptidase Caspase-2, helps to implement a CM-specific differentiation program that limits ploidy during postnatal heart development. DNA content analyses show that PIDDosome-loss causes a cell-autonomous increase in nuclear and cellular CM ploidy. Remarkably, increased ploidy does not affect cardiac structure nor function. PIDDosome-imposed ploidy restriction commences at postnatal day 7 (P7), reaching a plateau on P14. PIDDosome activation requires ANKRD26, targeting PIDD1 to mother centrioles. Opposite to prior observations in liver development, the PIDDosome limits CM polyploidization in a p53-independent manner but reliant on *p21/Cdkn1a*, a notion supported by nuclear RNA sequencing and genetic deletion experiments. Our results provide new insights how proliferation of polyploid CMs is restricted during postnatal heart development.

## 1. Introduction

Ischemic heart disease (IHD) is one of the leading causes of mortality worldwide with nearly 9 million deaths documented in 2017^1^. The primary cause of a reduced heart function is the initial loss of cardiomyocytes (CMs) and the inability of humans to compensate it^2,3^. In order to reverse this loss, induction of proliferation in pre-existing CMs appears to be a promising approach but is still not applicable due to a clear gap of knowledge of the terminal differentiation process. The terminal differentiation program is activated postnatally and is characterized by several tightly regulated steps in order to generate mature and functional CMs. After birth, CMs undergo a decrease of cell cycle activity, which is coupled with a change of metabolism from glycolysis to fatty acid oxidation, and an increased maturation status of the sarcomere apparatus^4,5^. Within the first postnatal week in rodents, the majority of CMs fail cytokinesis during mitosis resulting in binucleated CMs^2,6^. In the second to third postnatal week, a last wave of DNA synthesis occurs in a small subpopulation of binucleated CMs re-entering the cell cycle and thus, increasing nuclear ploidy^7^. Consequently, the adult mouse and rat heart consist of ∼80 to 90% binucleated CMs, while the adult human heart contains an estimated ∼25% binucleated and ∼25 to 40% polyploid mononucleated CMs^2,8–10^. Notably, recent studies showed that polyploidy limits cardiac regeneration by reducing the proliferation capacity of CMs^11,12^. Therefore, a better understanding of mechanisms controlling CM polyploidization is of great interest to uncover new avenues to treat heart diseases based on the induction of CM proliferation.

A common feature of mono- and binucleated polyploid cells is the presence of supernumerary centrosomes^13^. The centrosome is formed by two centrioles, referred to as mother and daughter, respectively, encapsulated in a pericentriolar matrix and acts as the major microtubule organizing center (MTOC) in most animal cells and is best known for its role in forming the mitotic spindle poles^14^. Mother centrioles differ from their daughters because they recruit distal appendage proteins, including ODF2, CEP83, SCLT1 and ANKRD26^15^. Postnatal binucleated CMs show extra centrosomes. In fact, ∼ 65% of postnatal day 3 (P3) binucleated CMs contain 4 centrioles, while ∼ 27% have 3 centrioles, with 2 mother centrioles^16^. Moreover, once stimulated with pro-proliferative factors *in vitro*, P3 binucleated CM can re-enter the cell cycle resulting in the formation of either a binucleated cell or two mononucleated polyploid cells ^16^. This shows that polyploid CMs have the intrinsic capability to cycle, but efforts to induce this process in adult CMs have largely failed due to the lack of knowledge of mechanisms controlling polyploidization-related cell cycle exit.

Polyploid cells with extra centrosomes can undergo a p53-dependent cell cycle arrest or cell death response ^17,18^. This is initiated by the “PIDDosome”^19^, a multiprotein complex formed by the auto-processed form of PIDD1 (P53-Induced death domain protein 1), termed PIDD1-CC, which recruits the bipartite adapter RAIDD/CRADD (RIP-Associated ICH1/CED3-homologous protein with Death Domain), and the pro-form of Caspase-2, a member of the family of cysteine-driven proteases known to control cell death and inflammation^20^. PIDD1 localization to the mother centrioles via the distal appendage protein ANKRD26, as well as centrosome clustering are requirements for the activation of this signaling complex, leading to Caspase-2 activation^21,22^. Active Caspase-2 proteolytically inactivates MDM2, the master regulator of cellular p53 levels, and thereby stabilizes p53 protein to promote a p21-dependent cell cycle arrest^19^.

Although first described in cancer cells forced to fail cytokinesis^19^, absence of either Caspase-2, RAIDD, or PIDD1, similar to p53- or p21-loss itself, increases ploidy in murine primary hepatocytes as part of a physiological polyploidization program during liver organogenesis^23^. Interestingly, the same study has shown that this function of the PIDDosome is preserved during liver regeneration, and likely also acts in human liver regeneration^23^. These findings raise the question of whether the PIDDosome and p53/p21 are also involved in controlling the scheduled polyploidization of CMs during postnatal heart development.

## 2. Methods

### Animals

Generation and genotyping of *Casp2^−/−^*, *Casp2*^fl/fl^, *Raidd^−/−^*, *Pidd1^−/−^mT*/*mG*, *p53^−/−^, p73^−/−^* and *p21^−/−^* mice was previously described^24–30^. *XMLC2-Cre* (*XMLC2^+^*) mice were a kind gift from Prof. Felix Engel^31^. In order to generate *XMLC2^+^Casp2^fl/fl^mice*, *XMLC2^+^* mice were crossed with the *Casp2^fl/fl^*mice ^32^. *XMLC2^+^*mice were crossed with a switchable Tomato and GFP reporter allele (mT/mG) in the Rosa26 locus, allowing expression of membrane bound versions of both fluorescent proteins. In order to generate *Casp2/p21^−/−^* double knockout (dKO) mice, *Casp2^−/−^* mice were crossed with *p21^−/−^* mice. All strains used were maintained on a C57BL/6N background and housed at the Animal Facility of the Medical University of Innsbruck under specific pathogen free conditions. *Ankrd26^−/−^*mice were generated, maintained and genotyped as previously described^33^ and heart tissues from littermates were kindly provided by Prof. Andrew J. Holland. In all experiments, age-matched animals were used indiscriminate of sex.

### Adult and postnatal cardiomyocyte isolation

Adult hearts from 3-month-old animals of the indicated genotypes were sacrificed by CO2 asphyxiation and cervical dislocation according to the governmental and international guidelines on animal experimentation. Adult cardiomyocytes were isolated by following the protocol of a Langendorff-free method published by Ackers-Johnson and colleagues^34^ with minor changes. To the Collagenase buffer, 30 µM CaCl_2_ was freshly added to optimize the digestion step. In order to have a final concentration of 0.5 mg/ml, Collagenase II (255 units/mg, 17101-015, Life Technologies, USA) and Collagenase IV (310 units/mg, 17104-019, Life Technologies) were used at 16.52 mg and 16.81 mg, respectively, as the unit concentrations of these enzymes were different from the original protocol. Moreover, the freshly made collagenase buffer, was pre-warmed on a heating magnetic stirrer at 37°C just before being injected in the left ventricle (LV). As first step, 7 ml of EDTA buffer was injected into the apex of the right ventricle with 1ml/min speed as described in the original protocol. Then, 20 ml EDTA buffer was injected into the LV over 10 minutes, and subsequently 3 ml of Perfusion buffer was injected into the LV with 1 ml/min speed. 40 ml of Collagenase buffer was finally injected into the LV over 13 minutes. However, after 8 minutes the heart was examined to see whether it has lost its color and structure and if it appeared over-digested. If this was the case, the digestion step was stopped at 8 minutes. After heart digestion and creating ∼1 mm^3^ pieces, the tissue trituration was done by gently pipetting for 2 minutes, followed by the addition of Stop buffer and further gentle pipetting for another 4 minutes. Adult cardiomyocyte vitality was checked after each isolation via Trypan blue staining. In order to assess cellular ploidy, cells were then centrifuged for 2 min at 100 x g, fixed in 4%PFA diluted in PBS for 20 mins at RT and then washed 3 times with PBS.

Hearts from postnatal day 1 (P1), P7 and P10 mice from the C57BL/6N mouse strain were dissected upon decapitation with operating scissors (RS-6845, ROBOZ, USA), the atria were removed, and the remaining ventricles were minced. For the isolation of postnatal cardiomyocytes, mouse hearts were isolated and digested utilizing the gentleMACS Dissociation kit (130-098-373, Milteny Biotech GmbH, Germany) according to the manufacturer’s instructions. For cardiomyocyte enrichment, cells were pre-plated in DMEM-F12/Glutamax TM-I (10565, Life Technologies)/10% fetal bovine serum (FBS, F0804, Sigma-Aldrich, USA)/penicillin (100 U/ml)/streptomycin (100 µg/ml) (Pen/Strep, P0781, Sigma-Aldrich). After 1.5 h, non-attached cells, enriched in cardiomyocytes, were collected, centrifuged for 10 min at 300 x g, resuspended in DMEM-F12, Glutamax TM-I containing 3 mM Na-pyruvate, 0.2% BSA, 0.1 mM ascorbic acid, 0.5% Insulin-Transferrin-Selenium (100x, 41400045, Life Technologies), 1% FBS, penicillin/streptomycin (100 U/mg/ml) and counted. Postnatal cardiomyocytes were washed once with PBS (14190169, Thermo Fisher Scientific, USA) and the cell pellet was snap-frozen in liquid nitrogen to preserve RNAs.

### Cardiomyocyte nuclei isolation and flow cytometric analyses

The isolation of CM nuclei was performed by following the previously described protocol^35,36^ with minor changes. Snap-frozen ventricles from selected genotypes were cut in 1 mm^3^ pieces and transferred into a falcon containing 15 ml of lysis buffer and then, homogenized with a TP 18/10 Ultra-Turrax probe homogenizer (IKA, Germany) at 20 000 rpm for 20s. In order to isolate the CM nuclei, the 15 ml lysis buffer solution with the homogenized tissues was passed 8 times up-and -down through a 20 ml syringe with a 20 G needle. The crude nuclei isolate was filtered using at first, a 100 µm cell strainer, and subsequently, a 70 µm cell strainer. After centrifugation for 10 min at 700 x g at 4°C, the nuclei pellet was dissolved in 5 ml sucrose buffer and then, topped up to 20 ml total solution. 10 ml sucrose buffer was added into a 1% BSA/PBS pre-coated ultra-centrifuge tube and was overlayed by the 20 ml nuclei suspension before spinning for 60 min at 13 000 x g at 4°C. After centrifugation, the sucrose buffer and any remaining liquid were quickly removed and the nuclei at the bottom of the tube were resuspended in 700 µl of nuclei storage buffer. Next, 600 µl of nuclei solution was incubated with a rabbit anti-PCM1 antibody (1:300, HPA023370, lot. number: 000007967, Sigma-Aldrich,) overnight at 4°C with constant shaking. 100 µl of the nuclei solution was used for the negative control of the staining. On the next day, both nuclei samples were washed by adding 2 ml of PBS, centrifuged for 10 min at 700 x g at 4°C and then, the nuclei pellets were incubated with goat anti-rabbit Alexa 647-conjugated antibodies (1:500, Life Technologies) for 60 min at 4°C. After 1 hour, the samples were washed with PBS and incubated with Propidium Iodide (1:25 of a 1 mg/ml solution). The gating strategy for ploidy analysis is shown in Suppl. Figure 1A, nuclear DNA content was measured on a flow cytometer (LSR-Fortessa, BD Biosystems, USA) and data were analyzed quantitatively, excluding doublets, using FlowJo (FlowJo X, LLC).

### Immunohistology

In order to have cardiomyocytes in a relax status (diastole), hearts were injected with a cardioplegic solution (25 mmol/L KCl) in the LV. Once the hearts stopped beating, they were cut from the aorta branch and placed in 4% PFA diluted in PBS overnight at RT. On the next day the fixed hearts were dehydrated and paraffinized. For hematoxylin and eosin staining, 4 µm frontal heart sections were deparaffinized, and rehydrated using graded xylol and ethanol incubation steps and then stained. At least 6 different hearts per genotype were analyzed. After the staining, sections were mounted and images acquired on NanoZoomer S210 Digital slide scanner (C13239-01, Hamamatsu photonics, Japan).

### Cryosections

Hearts from mentioned genotypes were pre-fixed in 4% PFA overnight at 4°C and on the next day, washed 3 times with PBS and incubated in 15% sucrose diluted in MilliQ water for 4h at 4°C. After the 15% Sucrose solution, hearts were submerged in 30% sucrose diluted in MilliQ water overnight at 4°C. Afterwards, hearts were embedded in Tissue-Tek O.C.T. compound tissue-freezing medium (4583, Sekura, Germany), frozen in liquid nitrogen, and sectioned with a Microm HM550 (Thermo Fischer Scientific) (10 µm). Once the tissues were cut, the sections were left to better adhere on the slide for 30 min at RT.

### Immunofluorescence analyses

Immunostainings were performed as previously described^37^, but after washing and incubation in fresh staining solution without antibodies, the CM suspensions were centrifuged for 2 min at 100 x g at RT. Before antibody incubation, freshly isolated and PFA-fixed CMs were permeabilized with 0.5% Triton X-100/PBS for 10 min at RT. Primary antibodies: rabbit anti-Pan cadherin (1:100, C3678, Sigma-Aldrich,), mouse anti-sarcomeric-µ-actinin (1:200, ab9465, Abcam). After overnight incubation, primary antibodies were detected by using Alexa Fluor™ 488, Alexa Fluor™ 594 or Alexa Fluor™ 647 conjugated antibodies (1:500, Life Technologies, USA). DNA was visualized with SYTOX™ Green Nucleic Acid Stain (1:200,000 in PBS with 0.1% triton and 10% horse serum, S7020, Thermo Fisher Scientific). Samples were diluted in PBS in order to acquire pictures. Confocal images were captured on a spinning disk confocal laser scanning microscope (Cell Voyager CV1000, Yokogawa, Tokyo, Japan).

FFPE heart sections were boiled in antigen retrieval buffer (1 mM EDTA, pH 8.0) for 8 min in a microwave (700W). After the antigen retrieval step, slides were kept for 30 min at RT and rinsed 3 times in PBS. Next, the slides were incubated in WGA (1:100 diluted in PBS, W11261, Thermo Fisher Scientific,) for 10 min at RT and then, washed 2 times with PBS. After this step, immunostainings were performed as previously described^37^. As primary antibody a goat anti-Troponin I (1:500, ab56357, Abcam) was used, which was detected by an Alexa Fluor™ 594 conjugated antibody (1:500, Life Technologies, USA). DNA was visualized with 0.5 µg/ml DAPI (4′,6′-diamidino-2-phenylindole) (Sigma Aldrich). Samples were mounted with Fluoromount-G™ (Thermo Fisher Scientific). Images were captured on a Zeiss Axiovert 200 M microscope using the VisiView 4.1.0.3 (Visitron Systems) acquisition software.

Immunostainings of pre-fixed heart cryosections were permeabilized with 0.5% Triton in PBS for 10 min at RT. Subsequently, the above-mentioned immunostaining protocol^37^ was used with a minor modification of the time of secondary antibody incubation, set to 1h at RT. High resolution images were captured on a LSM980 confocal laser scanning microscope (ZEISS, Germany).

### RNA isolation and qRT-PCR

Postnatal CM pellets were processed in order to extract total RNA using TRIzol TM Reagent (15596018, Invitrogen, USA) according to the manufacturer. 1µg of total RNA was used for generation of cDNA (iScript cDNA synthesis kit, 170–8891, BioRad, USA). For quantitative real-time PCR (RT-qPCR) experiments 140 ng of cDNA per each reaction was used. RT-qPCR assays were performed in experimental triplicates for each biological replicate using Luna® Universal Probe One-Step RT-qPCR Kit (E3006, New England biolabs, USA) in a StepOne Plus real time PCR system (Applied Biosystems, USA). Relative gene expression was calculated based on ΔCt values using *mGapdh* as housekeeping gene.

### Bulk nuclear RNA sequencing (RNAseq) pre-processing and analysis

For extraction of nuclear CM RNA, postnatal cardiomyocyte nuclei were isolated and stained using anti-PCM1, as described above. Subsequently, cardiomyocyte nuclei were sorted based on their PCM1 staining by using a BD FACSAria™ III Cell Sorter (648282, BD Bioscience). Importantly, in order to obtain enough RNA material from the isolated cardiomyocyte nuclei, several hearts from the same postnatal days were pooled together per biological replicate (P1: 15 hearts, P7: 6 hearts, P14: 2 hearts). Afterwards, the low-binding Eppendorf tubes containing the sorted cardiomyocyte nuclei were centrifuged for 10 min at 700 x g at 4°C. The resulting nuclei pellets were processed for RNA extraction by utilizing the Quick-RNA MicroPrep Kit (R1050, Zymo Research, USA) following the manufacturer’s protocol. RNA-sequencing library were prepared by Lexogen NGS Services (Vienna, Austria) using the QuantSeq 3′ mRNA-Seq Library Prep Kit FWD for Illumina and following the low-input protocol. Sequencing was performed on an Illumina NextSeq 2000 at Lexogen NGS Services to produce 100bp single-end reads for each sample. Raw RNA sequencing reads were quality-controlled with FastQC (v0.11.8)^38^ and preprocessed with cutadapt (v4.0)^39^ to trim poly-G stretches resembling sequencing artefacts, trim low-quality bases from the 3’end, trim adapter and poly-A sequences introduced by the sequencing strategy, remove low-quality reads (more than 1 expected error and/or more than 30% N-bases) and remove short reads (less than 20 nucleotides). Processed reads were aligned against the mouse reference genome GRCm39 from Ensembl (v108)^40^ using STAR (v2.6.1e)^41^. All following analyses were performed within R v. 4.2.1. The number of reads per gene (considering the full gene, i.e. both exonic and intronic sequences) was counted with HTSeq (v2.0.3)^42^. All following analyses were performed within R v4.2.1. Gene count normalization and differential gene expression analysis were performed with the R package limma (v. 3.52.4)^43^. Genes were considered as significantly differentially expressed with an adjusted p-value < 0.05 and an absolute log2 fold change > 1. Gene set enrichment analysis was carried out using the R package gage (v. 2.46.1)^44^, based on a selection of gene sets retrieved with the R package msigdbr (v. 7.5.1)^45^ (included gene set collections: hallmark, canonical pathways excluding WikiPathways, transcription factor targets, Gene Ontology). Gene sets were considered as differentially expressed when presenting with a plain p-value < 0.05.

Before gene count normalization, differential gene expression analysis, and gene set enrichment analysis, three samples (P1 *XMLC2^-^Casp2^fl/fl^*replicate 1, P14 *XMLC2^-^Casp2^fl/fl^* replicate 3, P14 *XMLC^+^Casp2^fl/fl^* replicate 1) were excluded due to low quality. The number of detected genes (at least one read count) was particularly low for P14 *XMLC2^-^Casp2^fl/fl^*replicate 3 (1676 genes) and P14 *XMLC^+^Casp2^fl/fl^* replicate 1 (3893 genes) compared to the other samples (between 7223 and 17057 genes) (data not shown). P1 *XMLC2^-^Casp2^fl/fl^* replicate 1 clustered particularly far away in a Principal component analysis (PCA) of all samples (PCA calculated with R package labdsv v2.1-0^46^ (Suppl. Figure 3A). A fuzzy clustering of all samples showed clusters of genes (clusters 6 and 7) with high expression level in P1 *XMLC2^-^Casp2^fl/fl^* replicate 1 compared to replicates 2, 3, and 4 (fuzzy clustering performed with R package Mfuzz v2.56.0^47^ (Suppl. Figure 4). The clustering was calculated on the gene log2 fold changes with respect to the average gene counts of the four P1 *XMLC2^-^Casp2^fl/fl^*replicates, after each sample’s raw counts were normalized with the average count of its interquartile range count values. Subsequent functional enrichment analysis of gene cluster 6 and cluster 7 returned significantly enriched gene sets (adjusted p-value < 0.05) associated with mitochondria, ribosomes and translation (Suppl. Figure 3B). Functional enrichment was calculated with Fisher’s exact test, based on the same gene set collection as described above. As these results are indicative of a cytoplasmic contamination of P1 *XMLC2^-^Casp2^fl/fl^* replicate 1, this sample was not considered for further analysis. <u>GEO submission number</u> (GSE275946). Reviewer token (uvyhckkqxvwfvap).

### Tissue lysis and SP3 proteome clean up

Pulverized tissues from P7 hearts of *XMLC2^-^Casp2^fl/fl^* and *XMLC^+^Casp2^fl/fl^* mice were lysed, reduced, and alkylated by heating for 10 min at 95 °C under shaking at 1500 rpm in lysis buffer containing 6 M guanidine hydrochloride (GuHCl, Sigma), 10 mM tris(2-carboxyethyl) phosphinehydrochloride (TCEP, Thermo Fisher), 40LmM chloroacetamide (CAA, Sigma), and 200 mM 3-[4-(2-Hydroxyethyl)piperazin-1-yl]propane-1-sulfonic acid (HEPPS, Sigma), pH 8.5. Protein concentration was estimated using a BCA assay (Thermo Fisher). For each sample, 200 μg of protein were incubated with 10 μL of SP3 beads ^48^ (1:1 mix of Sera-Mag Speed Beads A and B, Cytiva). Pure ethanol (EtOH, VWR) was added to achieve a final concentration of 80% (v/v), and samples were incubated at room temperature in a thermoshaker for 18 min at 800 rpm. The samples were then placed on a magnetic rack for 2 minutes to immobilize the SP3 beads, after which the supernatant was discarded. The beads were washed twice with 1 mL of 80% (v/v) EtOH in water and once with 800 µL of acetonitrile (ACN, VWR).

### TMT labeling and protein digestion

SP3-bound proteins were resolubilized with 120 μL of labeling buffer containing 2 M GuHCl, 10LmM TCEP, and 200LmM HEPPS, pH 8.5. The samples were incubated at room temperature in a thermoshaker for 30 minutes at 800 rpm, with 1 minute sonication in a water bath every 10 minutes. TMT10-plex labeling (Thermo Fisher) was performed as previously described^50^. Briefly, 0.8 mg of TMT reagent dissolved in 30 µL of 100% anhydrous ACN (Sigma) was added to the samples to label free N-termini (protein N-termini and lysine residues). The labeling reaction proceeded for 1 hour at 20 °C in a thermoshaker at 400 rpm. The reaction was quenched by adding 5 µL of 1 M Tris/HCl pH 8.5 (Sigma) and incubating for 30 min at 25 °C. After quenching, the samples were combined, and SP3 cleanup was repeated to remove excess reactants as described above. The combined proteome was resuspended in 100 mM 4-(2-hydroxyethyl)-1-piperazineethanesulfonic acid (HEPES, Sigma), pH 7.6, and digested overnight at 37 °C with trypsin at 1:50 protease:protein ratio (w/w).

### N-terminome negative selection

To remove internal trypsin-generated peptides, the digest was incubated overnight with 30 mM sodium cyanoborohydride (Sigma) and the amine-reactive hyperbranched aldehyde-derivatized polymer^52^ (HPG-ALDII https://ubc.flintbox.com/technologies/888fc51c-36c0-40dc-a5c9-0f176ba68293) at 1:5 peptide:polymer ratio (w/w). The reaction was quenched by adding Tris/HCl pH 7.6 to a final concentration of 100 mM. The resulting sample was centrifuged using an Amicon 30 kDa filter (Sigma) to collect TMT-labeled N-termini in the flow through. The TAILS sample was acidified to 1% formic acid (FA, Carlo Erba), desalted and further fractionated into six fractions using the high pH RP fractionation^54^,employing a self-packed StageTip with five disks of C18 material (3LM Empore). The peptides were eluted with 25 mM ammonium formate pHL10 (Sigma), using increasing concentrations of ACN (5, 7.5, 10, 12.5, 15, 17.5, and 50%). Finally, the seven fractions with flow-through were combined into six fractions (5L+L50%, 7.5%, 10%, 12.5%, 15%, and 17.50%L+Lflow-through), the peptides were dried in vacuo and stored at −20 °C until further use.

### nLC-MS/MS

Nano flow LC-MS/MS measurements were performed using a Dionex Ultimate 3000 UHPLC+ system coupled with an Orbitrap Eclipse mass spectrometer (Thermo Fisher). Peptides were delivered to a trap column (75 μm i.d. × 2 cm, packed in-house with 5 μm Reprosil C18 beads, Dr. Maisch) and washed using 0.1% FA at a flow rate of 5 μL/min for 10 min. Subsequently, peptides were transferred to an analytical column (75 μm i.d. × 40 cm, packed in-house with 1.9 μm Reprosil C18 beads, Dr. Maisch) at a flow rate of 300 nL/min. Peptides were chromatographically separated using an 80 min linear gradient from 8 to 34 % of solvent B (0.1% FA, 5% DMSO (Sigma) in ACN and solvent A (0.1% FA, 5% DMSO in water).

The Orbitrap Eclipse was operated in a data-dependent acquisition (DDA) to automatically switch between MS and MS/MS. Briefly, survey full-scan MS spectra were recorded in the Orbitrap from m/z 360 to 1500 at a resolution of 60K, using an automatic gain control (AGC) target value of 100% and maximum injection time (maxIT) of 50 ms. For the MS3-based TMT method, initial MS2 spectra for peptide identification were recorded in the Orbitrap at a resolution of 15K with a top speed approach using a 3-s duration (isolation window m/z 0.7, AGC target value of 100%, maxIT of 22 ms). Fragmentation was set to HCD, with a NCE of 34%. Then, for each peptide precursor, an additional MS3 spectrum for TMT quantification was obtained in the Orbitrap at 30K resolution with enhanced resolution mode enabled (AGC of 500%, maxIT of 54 ms). The precursor was fragmented as for the MS2 analysis, followed by synchronous selection of the 10 most intense peptide fragments and further fragmentation via HCD using a NCE of 55%. Dynamic exclusion was set to 90 s.

### Data analysis of N-terminomics

Raw mass spectrometry data were processed using the FragPipe software (version 21.1) with its built-in search engine MSFragger version 4.0^56^. Spectra were searched against the mouse UniProtKB database UP00000589 (63,300 entries including isoforms, downloaded on 04.2024). Default parameters for a TMT10-MS3 search were employed, with a defined precursor tolerance of 20 ppm and enzyme semi-tryptic specificity for database digest set to trypsin_r. Methionine oxidation (+15.9949) and protein n-termini acetylation (+42.0106) were set to variable modifications. Cysteine carbamidomethylation (+57.02146), and TMT (+229.16293) on lysine and peptide n-termini were added as static modifications.

After peptide-to-spectrum matches (PSM) rescoring via percolator^58^, identifications were adjusted to 1% false discovery rate (FDR) at the protein, peptide and PSM levels. TMT integrator, included in FragPipe, was used to perform MS3-based quantification of the detected peptide features, using default settings.

Data analysis was performed with the Perseus software (version 2.0.10.0.)^60^. Peptide identifications were filtered to remove contaminants before performing data normalization of the log2-transformed TMT intensity values by median centering, as implemented in Perseus. For statistical analysis, only peptides that had been quantified in at least 3 biological replicates were retained, and missing values were imputed from the normal distribution in Perseus, using default parameters. Potential caspase-2 substrates enriched in the WT over the KO samples were identified using the Significance A test^62^ on the distribution of TMT ratios, using a Benjamini-Hochberg FDR threshold of 5%.

The mass spectrometry proteomics data have been deposited in the ProteomeXchange Consortium via the PRIDE partner repository^64^ with the dataset identifier PXD060421 (Website: http://www.ebi.ac.uk/pride, Username, Password: BHrAjVHxayye).

### Imaris image analysis

Nuclear and cellular ploidy measurements were performed by following the indications published by Bensley and colleagues^10^. More in detail, firstly, reconstituting the individual nuclei by the 3D volume module (“3D View” in Imaris) was accomplished and then, analyzing the mean intensity of the Sytox Green was achieved by using the “surface” Imaris software package. Importantly, since signal intensity is fundamental for accurate ploidy measurement, the same setting of laser power, voltage, offset, and pinhole across the board was kept constant for all the experiments. Single nuclei from binucleated CMs were taken as reference for diploid nuclei in this analysis. For nuclear and cellular ploidy analysis, images were captured on a spinning disk confocal microscope, CV1000 Cell voyager (Yokogawa, Japan).

### Statistical analysis

Data are expressed as the mean ± standard deviation (SD) or standard error of the mean (SEM) of at least three independent experiments if not stated otherwise. Statistical significance of differences was evaluated by either Student’s t test, or 1way or 2way ANOVA followed by Bonferroni’s post-hoc test, Sidak’s or Tukey’s multiple comparisons test (GraphPad Prism 9.0). p < 0.05 was considered as statistically significant.

## 3. Results

### The PIDDosome controls cardiomyocyte ploidy

In order to decipher the role of the PIDDosome during CM polyploidization, we have quantified the nuclear ploidy levels of freshly-isolated CMs from 3-month-old mice lacking individual PIDDosome components (Figure 1A). Ploidy of single CM nuclei was assessed by 3D volumetric analysis of Sytox Green fluorescence intensity and correlated with the cellular ploidy status. Moreover, the average nuclear staining intensity of single nuclei from binucleated CMs was used to define 2N ploidy, since more than 90 % of binucleated CMs have diploid nuclei^10^. This analysis revealed that CM nuclei of binucleated cells from all three different PIDDosome knockout strains contain a higher nuclear ploidy, compared to nuclei from binucleated CMs from WT hearts. This indicates that binucleated CMs in *Pidd1, Raidd* and *Casp2-*deficient animals harbor an increased number of nuclei with DNA content > 2N (Figure 1B). To determine the exact nuclear ploidy of single CM nuclei from PIDDosome-mutant animals, we used a flow cytometry-based strategy. To discriminate CM nuclei from non-myocyte nuclei, we exploited the translocation of the pericentriolar matrix protein PCM1 from the centrosome to the nuclear envelope during terminal CM differentiation^49^ (Suppl. Figure 1A). Intriguingly, PIDDosome-mutant animals showed a more than two-fold increase in the percentage of tetraploid (4N) CM nuclei, compared to those isolated from WT animals (Figure 1C). Quantification of the cellular ploidy status of CMs showed that absence of the PIDDosome also causes a significant increase in multinucleated CMs (Figure 1D). Altogether, these results indicate that abrogation of the PIDDosome induces a pronounced increase in nuclear, as well as cellular polyploidization of CM. This suggests that in the absence of the PIDDosome, tetraploid binucleated CMs enter another round of cell cycle, characterized by cytokinesis failure generating either octoploid binucleated CMs or multinucleated CMs with diploid single CM nuclei.

**Figure 1.**
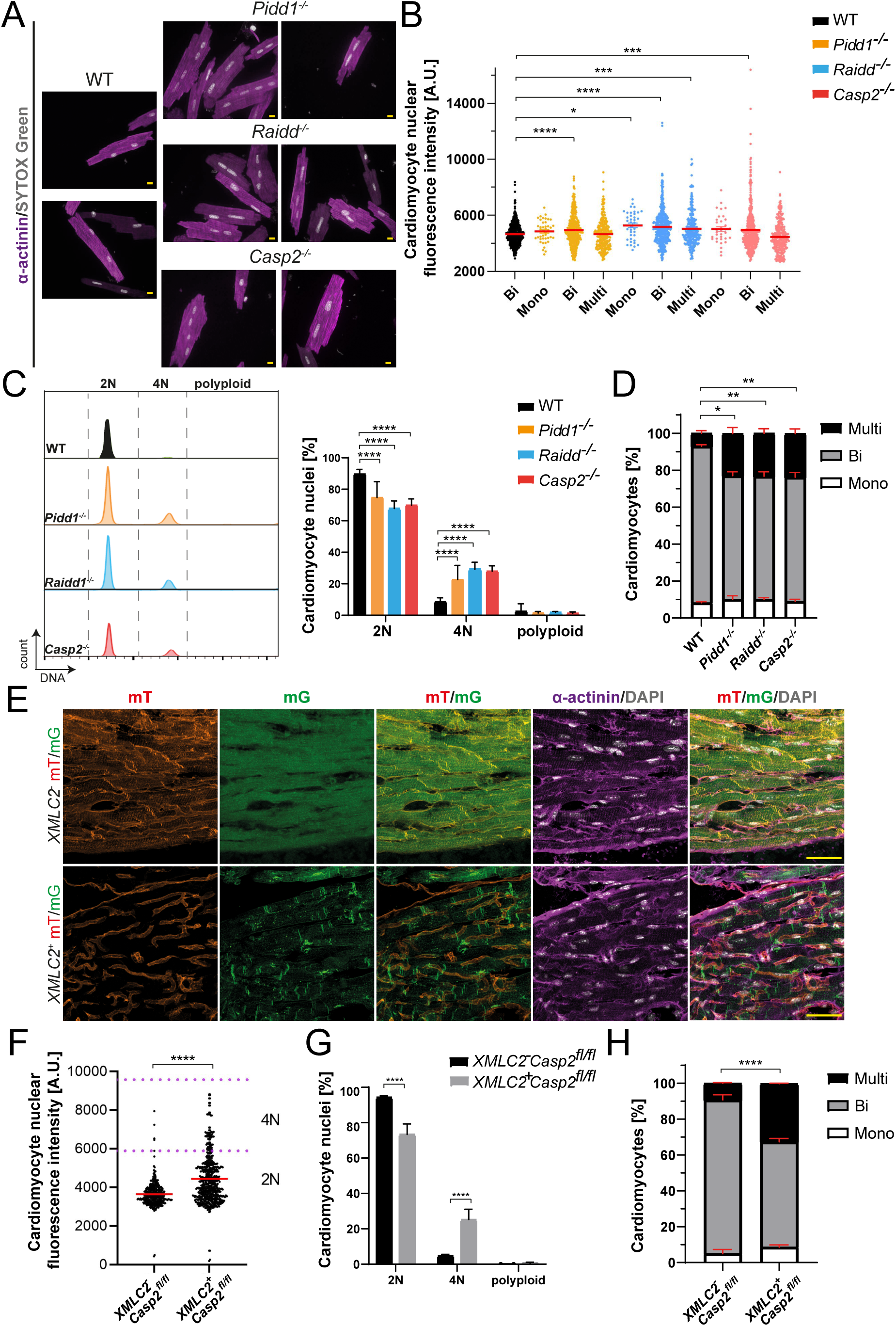
The PIDDosome regulates nuclear and cellular polyploidization of cardiomyocytes. (**A-B**) Ploidy measurements of single CM nuclei by 3D volumetric analysis of Sytox -green fluorescence intensity in correlation with the cellular ploidy status. **(A)** Representative pictures of freshly-isolated 3-month-old CMs from different genotypes. CMs were stained for the cardiac marker, α-sarcomeric actinin (purple), and a nuclear marker, Sytox-green (grey). Scale bars: 10 μm. **(B)** Scatter blots of quantification of Figure A. n: 4 hearts per genotype; 100 CMs per heart. (**C**) Representative histograms (left) and quantification (right) of the CM nuclear ploidy measured by flow cytometry in 3-month-old hearts from the indicated genotypes. n: 7 (WT), n:4 (*Pidd1^−/−^*), n:4 (*Raidd^−/−^*), n:5 (*Casp2^−/−^*).(**D**) Bar graphs of CM cellular ploidy of the indicated genotypes at 3-month-old. n: 4 hearts per genotype. 150 CMs per heart were quantified. Statistical analysis between the multinucleated group from the different genotypes is shown in the graph. (**E**) Validation of the cardio-specificity of the *XMLC2* promoter. Confocal pictures of cryosections of pre-fixed hearts stained for α-sarcomeric-actinin (purple) and DAPI (grey). Scale bars: 40 μm. (**F**) Quantification of CM nuclear ploidy by 3D volumetric analysis of Sytox green fluorescence intensity. n: 3 hearts per genotype. 100 CM nuclei per heart were analyzed. (**G**) Quantification of CM nuclear ploidy of the indicated genotypes in 3-month-old mice by flow cytometry. n: 4 (*XMLC2*^-^ Casp2^fl/fl^), n: 8 (*XMLC2*^+^ Casp2^fl/fl^). (**H**) Bar plots of CM cellular ploidy of the indicated genotypes at 3-month-old. n: 3 hearts per genotype; 150 CM nuclei per hearts were analyzed. Data are mean ± SEM (B,D,F,H) or ± SD (C,G) analyzed by One-way ANOVA (B) or Two-way ANOVA (C,D,G,H) or Two-tailed Student’s t test (F).* p < 0.05, **p < 0.01, ***p < 0.001 ****: p < 0.0001, n.s.: not significant.

To test if the observed phenotype in PIDDosome knockout mice is independent of external parameters, e.g., non-myocytes influencing postnatal CM polyploidization, we next generated a mouse mutant harboring a cardiac specific deletion of *Casp2* (named from now on, *XMLC2^+^Casp2^fl/fl^*) (Suppl. Figure 1B). To exclude that cardiac CRE expression under control of *XMLC2* promoter affects CM polyploidization and to confirm its cardiac-specificity, *XMLC2-Cre* (*XMLC2^+^*) mice were first crossed with a fluorescence reporter line, carrying a switchable Tomato/GFP reporter allele in the *Rosa26* locus, allowing expression of membrane bound versions of both fluorescent proteins (*mT/mG*). In the *mT/mG* model, all cells initially express the Tomato reporter prior to CRE recombination, but not GFP. CRE recombination leads to excision of both the *Tomato* cassette and the STOP signal, allowing expression of membrane-bound GFP (Suppl. Figure 1C). Consistently, upon *XMLC2*-driven CRE activation, CM membranes in heart cryo-sections were GFP-positive, while non-CM membranes remained Tomato-positive. Staining with α-sarcomeric actinin and DAPI was performed in addition to define cell identity and nuclear morphology, confirming specificity and effective CRE-recombination (Figure 1E). Importantly, measuring the CM nuclear ploidy in *XMLC2^+^* mice, excluded off-target effects on nuclear ploidy (Suppl. Figure 1D). 3D volumetric analysis of Sytox Green staining intensity, as well as flow cytometric analysis revealed that cardiac-specific *Casp2* deletion in *XMLC2^+^Casp2^fl/fl^* mice significantly increased the percentage of polyploid 4N CM nuclei, when compared to *Casp2^fl/fl^* control mice (*XMLC2^-^Casp2*^fl/fl^) (Figure 1F-G). In addition, the cellular ploidy of *XMLC2*^+^*Casp2^fl/fl^* CMs (Figure 1H) increased to a similar degree seen in *Casp2*^−/−^ animals (Figure 1D).

Taken together, these data indicate that the PIDDosome controls postnatal CM polyploidization, limiting not only the nuclear but also cellular ploidy in a cell autonomous manner.

### Increased CM ploidy does not affect cardiac structure or function

Since PIDDosome loss causes an increase of polyploid CMs in adulthood, we wondered whether this might affect the heart structure and consequentially, cardiac function. The heart ultra-structure of the different genotypes was analyzed by hematoxylin and eosin (H&E) staining on frontal sections of formalin-fixed and paraffin-embedded (FFPE) hearts. No obvious differences were observed in *Casp2^−/−^*, *Raidd^−/−^*or *Pidd1^−/−^* hearts compared to WT hearts (Figure 2A), nor in *XMLC2*^+^ *Casp2*^fl/fl^ *vs*. *XMLC2*^-^ *Casp2*^fl/fl^ hearts (Suppl. Figure 1E). Likewise, echocardiography of 7-10-weeks old mice lacking *Pidd1* confirmed that the observed increases in the polyploid CM population did not affect the cardiac function. In fact, there were no significant differences in ejection fraction (EF) and fractional shortening (FS) between *Pidd1^−/−^* and WT mice (Figure 2B-C).

**Figure 2.**
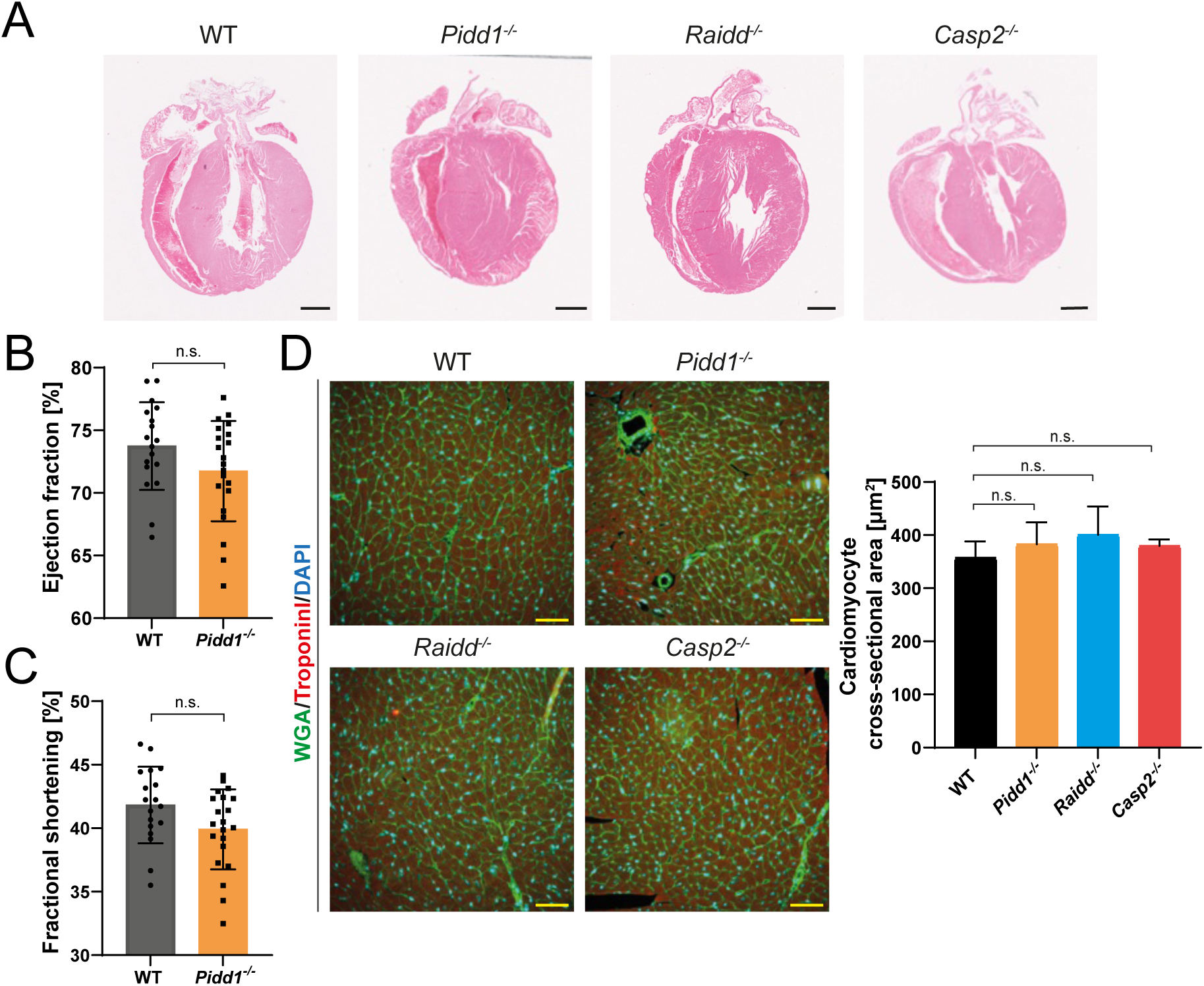
Heart structure and functionality are not influenced by the elevated PIDDosome-dependent CM ploidy. **(A)** Representative images of hematoxylin and eosin (H&E) stained FFPE heart sections of 3-month-old mice of the indicated genotypes. Scale bars: 1mm. (**B-C**) Quantification of the cardiac functions, ejection fraction (B) and fractional shortening (C), measured by echocardiography in 3-month-old mice. n: 21 (WT), n: 18 (*Pidd1*^−/−^). Statistical significance was assessed with Student’s t test. (**D**) Representative pictures (left) and cross-sectional area quantifications (right) of FFPE hearts from WT, *Pidd1*^−/−^, *Raidd*^−/−^ or *Casp2*^−/−^ mice stained for the membrane marker WGA (green), the cardiac marker Troponin I (red) and DAPI (blue) for the nuclei. Scale Bars: 50µm. n: 5 (WT, *Pidd1*^−/−^, *Raidd*^−/−^, *Casp2*^−/−^, ∼ 2100 CMs distributed within 7 pictures per heart were quantified). Data are mean ± SD, two-way ANOVA n.s.: not significant.

Polyploid cells have multiple copies of the DNA content, show increased nuclear size, and they are usually larger than their diploid counterparts^51^. Since ablation of PIDDosome components causes an increase in polyploid CM, we wanted to investigate whether this ploidy increase was associated with an increase of CM area. For this reason, frontal sections of (FFPE) hearts from *Casp2^−/−^*, *Raidd^−/−^, Pidd1^−/−^*and WT mice were stained with a plasma membrane marker, Wheat germ agglutinin (WGA), a cardiac marker, Troponin I, and DAPI. Quantification of the cross-sectional area of CMs, measured by a combination of WGA and Troponin I staining, revealed no differences across genotypes (Figure 2 D), suggesting that increases of ploidy are not associated with an increase in CM size, even though contrasting findings were made in the liver^53^.

Collectively, these data show that PIDDosome deletion and the subsequent increases in the number of polyploid CM do not alter the tissue architecture of the heart and more importantly, cardiac function in steady state.

### The PIDDosome restricts cardiomyocyte ploidy in early postnatal development

Mouse cardiomyocytes fail to undergo cytokinesis, thus becoming binucleated during the first week after birth^3,9^. As described by Alkass and colleagues^7^, a second wave of polyploidization occurs between the second and the third postnatal week. To define when the PIDDosome may be engaged during cardiomyocyte development, we first evaluated postnatal mRNA expression of the individual PIDDosome components by qRT-PCR from isolated CMs. All PIDDosome components showed low expression levels at P1, while at P7 mRNA levels of all components were found to be strongly increased. Interestingly, decreasing expression of these genes at P10 suggests that the PIDDosome components are under tight transcriptional control during postnatal heart development (Figure 3A). Next, in order to unveil when the PIDDosome exerts CM-specific ploidy control, hearts were isolated from WT as well as *Casp2^−/−^*, *Raidd^−/−^*or *Pidd1^−/−^* mice on P1, P7, P14 and P21 and CM nuclear ploidy was assessed by flow cytometry. As expected, on P1 no significant difference in the fraction of tetraploid CM nuclei was observed across genotypes (Figure 3B), consistent with diploid mononucleated CMs dominating at this developmental stage^3^. However, at P7 the population of tetraploid CM nuclei in PIDDosome mutant mice increased significantly, compared to the same population in WT mice (Figure 3C). This phenomenon coincides with the formation of binucleated CM at this time point during heart development^3^. Interestingly, the percentage of tetraploid CM nuclei reached a plateau at P14 as no further increases were noted on P21 (Figure 3D-E), matching numbers were seen in adult mice (Figure 1C). Flow cytometric ploidy analysis of *XMLC2^+^Casp2^fl/fl^* mice confirmed that this is a cell-autonomous effect (Figure 3F).

**Figure 3.**
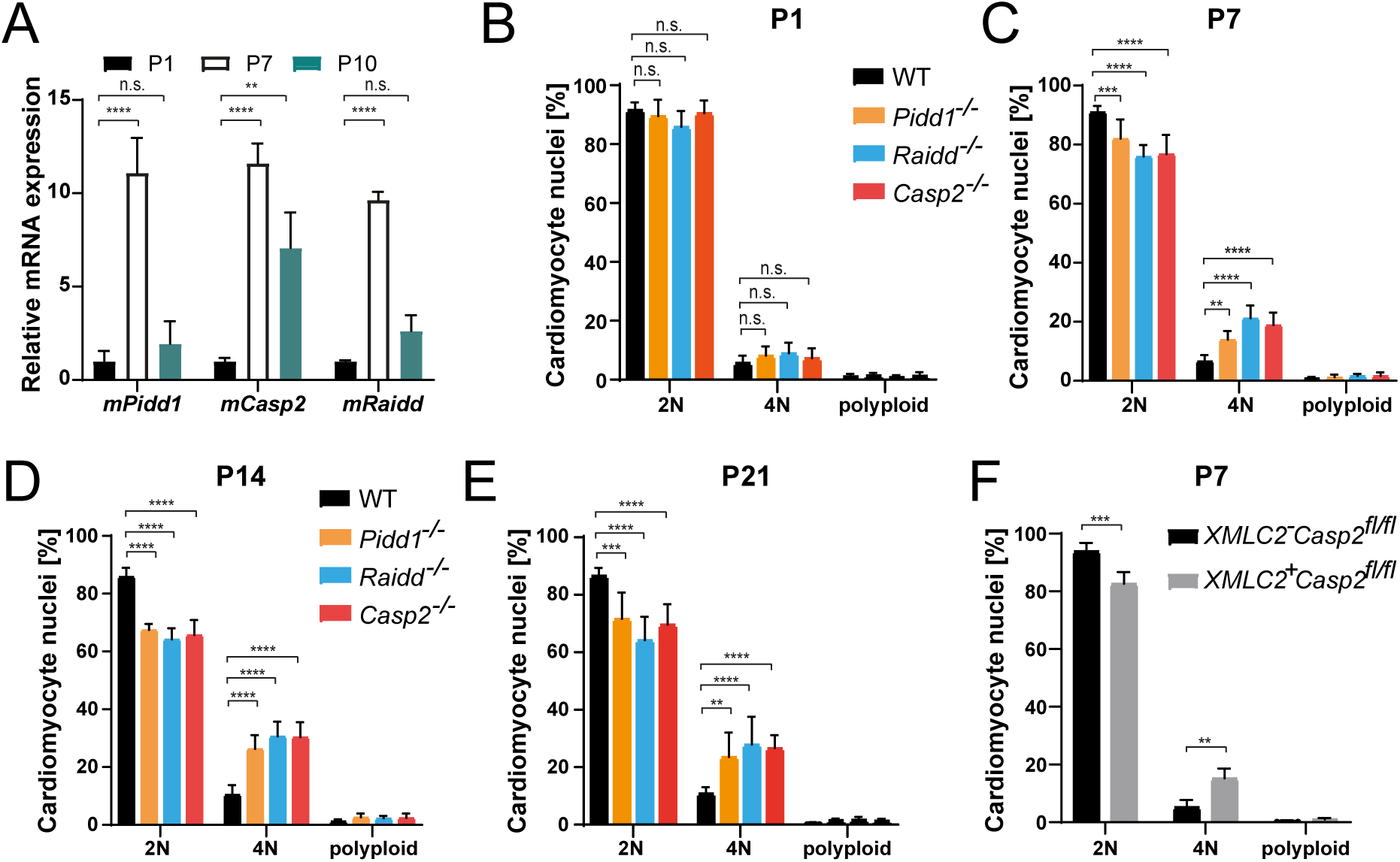
The PIDDosome controls CM ploidy levels starting on day P7 after birth. (**A**) PIDDosome mRNA levels were determined in isolated P1, P7 and P10 CMs from WT mice by qRT-PCR. n: 3 individual isolations per each postnatal timepoint. (**B**-**E**) Quantification of the CM nuclear ploidy measured by flow cytometry of P1 (B), P7 (C), P14 (D) and P21(E) from the indicated genotypes. n: 3-5 for P1 (5 hearts of each genotype per n), n: 5 for P7 (3 hearts of each genotype per n), n: 3 for P14 (2 hearts of each genotype per n), n: 4-6 for P21 (2 hearts of each genotype per n). (**F**) Quantification of the CM nuclear ploidy in *XMLC2^-^Casp2^fl/f^*^l^ and *XMLC2^+^Casp2^fl/fl^* mice at P7. n: 4 (*XMLC2^-^Casp2^fl/fl^*), n: 5 (*XMLC2^+^Casp2^fl/fl^*) (n: 3 hearts of each genotype per n). Data are mean ± SEM (A) or ± SD (B-F) analyzed by Two-way ANOVA. **p < 0.01, ***p < 0.001, ****p < 0.0001, n.s.: not significant.

Collectively, these data show that the PIDDosome controls CM polyploidization as early as P7.

### Extra centrosomes induce postnatal ploidy control in CM

PIDDosome activation depends on PIDD1 binding to extra mother centrioles via the centriolar distal appendage protein ANKRD26 in cancer cells^55^, as well as in primary hepatocytes^33^. Of importance, previous work has shown that P3 binucleated CMs contain 3 or 4 centrioles including extra mother centrioles ^16^ suggesting these may be involved in pathway activation. To evaluate the relationship of PIDDosome activation and centrosomes, we quantified the CM nuclear ploidy in hearts from 3-month-old mice lacking the molecule that links PIDD1 to mother centrioles, ANKRD26. Indeed, the percentage of tetraploid nuclei in *Ankrd26*^−/−^ mice was increased by two-fold, similar to what was observed in the PIDDosome knockout mice (Figure 4A).

**Figure 4.**
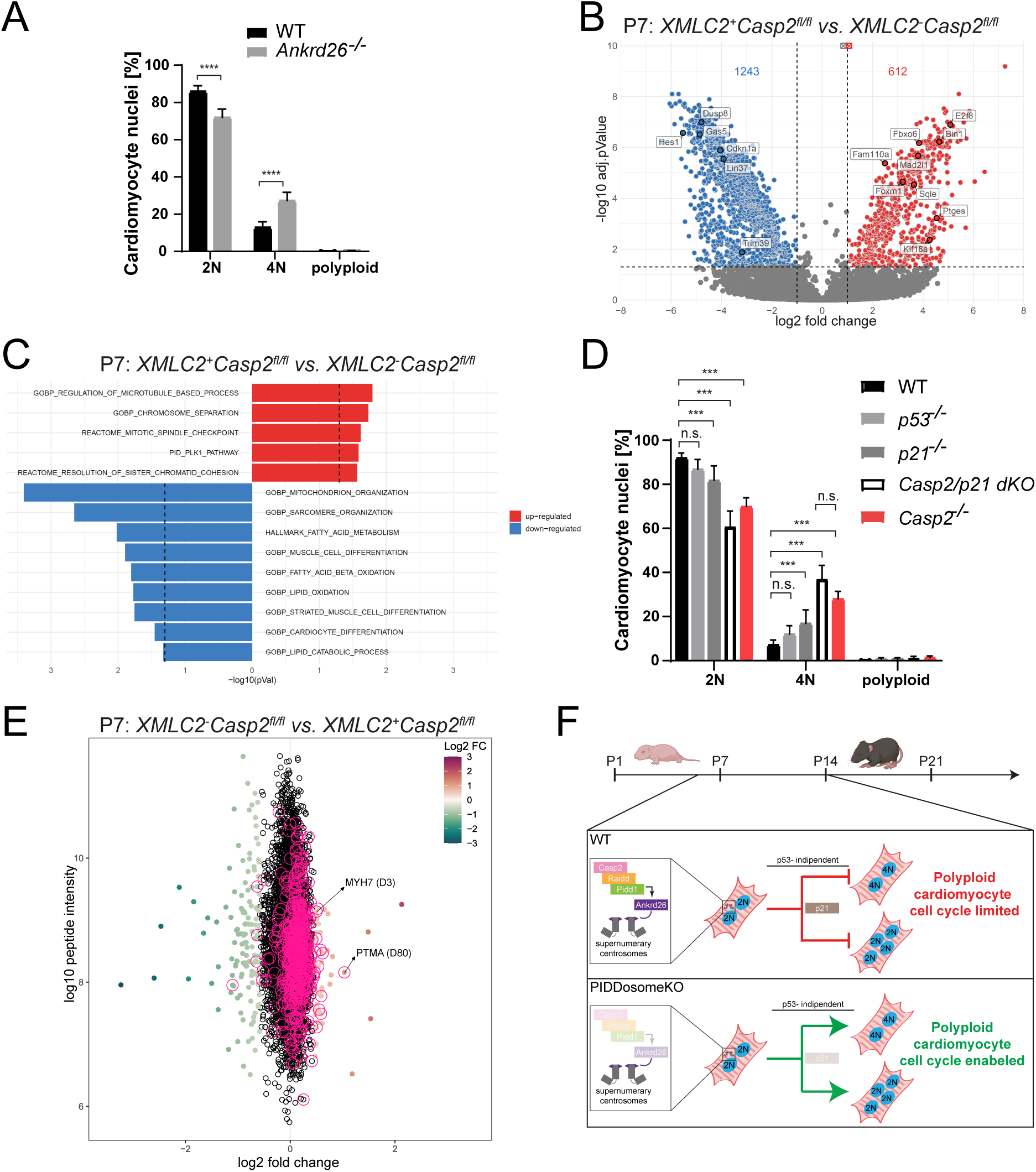
PIDDosome-mediated CM ploidy control depends on extra centrosomes but not p53. (**A**) Quantification of CM nuclear ploidy of 3-month-old *Ankrd26^−/−^* and WT (C57BL/6J) mice measured by flow cytometry. n: 4 per genotype. (**B**) Volcano plot of the differential gene expression analysis comparing *XMLC2^+^Casp2^fl/fl^* to *XMLC2^-^Casp2^fl/fl^* samples at P7. Dashed lines indicate an adj. p-value of 0.05 and a log2FC of −1 and 1. Numbers of up- or downregulated differentially expressed genes are indicated in the graph. (**C**) Selected significantly deregulated gene sets of the gene set enrichment analysis results comparing *XMLC^+^ Casp2^fl/fl^* to*. XMLC2^-^Casp2^fl/fl^* samples at P7. Dashed lines represent a plain p-value of 0.05. All data are mean ± SD analyzed by Two-way ANOVA. ****p < 0.0001, n.s.: not significant.(**D**) Bar plots of CM nuclear ploidy of the indicated genotypes. n: 10 (WT), n:7 (*p21*^−/−^), n:5 (*p53*^−/−^), n: 6 (*Casp2^−/−^/p21^−/−^*), n:5 (*Casp2*^−/−^, as in Fig. 1C) (**E**) Significance A plot showing the log2-fold change of N-termini measured by quantitative mass spectrometry in *XMLC2^-^Casp2^fl/fl^*vs. *XMLC^+^ Casp2^fl/fl^*heart tissues. Significantly up- or down-regulated n-termini (adjusted *p*-valueL<L0.05) are color-coded based on the observed fold-change. N-termini generated by cleavage C-terminal to aspartic residues are highlighted with a purple outline. (**F**) Cartoon summary: The PIDDosome exerts its CM ploidy controlling function within the first two postnatal weeks in mice. Activation depends on extra centrosomes but ploidy is regulated independently of p53 stabilization, yet engaging p21 downstream of Caspase-2 in a yet to be defined manner.

Our findings indicate that the supernumerary centrosome and mother centriole numbers in CMs trigger PIDDosome activation in an ANKRD26-dependent manner.

### Caspase-2-loss enhances expression signatures related to CM proliferation and maturation

In order to elucidate the biological processes through which the PIDDosome executes its ploidy-regulating function, P1, P7 and P14 CM nuclei were isolated from *XMLC2*^-^*Casp2^fl/fl^* and *XMLC2*^+^*Casp2^fl/fl^*mice and enriched by cell sorting based on PCM1 staining to perform bulk RNAseq (Suppl. Figure 2A). Gene set enrichment (GSEA) analyses supported the previously described developmental mRNA expression changes in CMs between P1, P7 and P14^57,59^. These included the noted upregulation of different ion channels in P14, compared to P7 specimens, a known feature of cardiomyocyte maturation^61,63^. In addition, gene sets associated with cell adhesion were upregulated in P1 compared to P7 CMs, as previously published^59^. Gene sets of RNA splicing, aerobic respiration and oxidative phosphorylation were also abundant in P7 CMs in both comparisons, P14 *vs.* P7 and P7 *vs.* P1 samples, as clear signs of cardiomyocyte maturation^65^. Interestingly, when comparing the nuclear transcriptomes of CMs isolated from P7 *vs.* P1, gene sets defining “mitochondrial protein complexes” and “mitochondrial organization” were also enriched, indicating that P7 CMs have a more mature mitochondria structure or organization compared to P1 CMs, confirming published results ^63,66^ (Suppl. Figure 2B-C). In addition, a trend towards the switching of genes which play a role in sarcomere structure and regulation, involved in the “Cardiac Fetal-to-Adult Contractile Isoform Transition”, was also confirmed between P1 and P7 CMs (Suppl. Figure 2D). This feature was previously associated with the developmental maturation of CMs^59,63^.

Since the PIDDosome starts to affect the CM ploidy status during the first week postnatal (Figure 3), we next investigated the differentially expressed genes in *Casp2*-depleted CMs (*XMLC2*^+^*Casp2^fl/fl^* mice), compared to CMs from control animals (*XMLC2*^-^*Casp2^fl/fl^* mice) at P7 (Figure 4B). Interestingly, genes related to CM proliferation (*Foxm1*^67–69^, *Gas5*^70,71^, *Sqle*^72^, *Dusp8*^73^), cell cycle regulation (*Cdkn1a*^74^, *Hes1*^75^, *Trim39*^76^, *Lin37*^77^, *Kif18a*^78,79^), ploidy control (*E2f8*^80^, *Mad2l1*^81^) and T-Tubule maturation (*Bin1*^82^) were found differentially expressed between genotypes (Figure 4C). Further, gene set enrichment analysis highlighted genes found in GO terms related to chromosome separation, mitotic spindle checkpoint, the PLK1 pathway, controlling also centrosome biogenesis, resolution of sister-chromatid cohesion and microtubule-based processes, were upregulated in *Casp2*-depleted CMs compared to controls, at P7. This suggests that Caspase-2-depleted CMs, which are already more polyploid compared to their control counterparts on P7, might experience mitotic errors and delays due to their increased DNA content. Importantly, downregulated GO terms in P7 *Casp2*-depleted CMs *vs.* P7 controls were associated with a reduction in fatty acid metabolism and muscle differentiation, as well as sarcomere organization (Figure 4C).

Together, this suggests that the absence of Caspase-2 in CMs causes delays in the naturally occurring terminal differentiation program, resulting in the presence of more immature and cell cycle active CMs, compared to control mice at P7.

### PIDDosome-driven CM ploidy control does not require *p53* but involves *p21*

Mechanistically, the PIDDosome regulates ploidy by the activation of the protease activity of Caspase-2 and subsequent MDM2 cleavage, leading to p53 activation. As this axis is conserved across several cell types^19,23^, we investigated whether it is also preserved in CMs. To this end, we have analyzed CM ploidy in 3-month-old *p53*^−/−^ mice. Surprisingly, the percentage of tetraploid CM nuclei in mice lacking this key ploidy regulator was not significantly different compared to that found in WT animals (Figure 4D). In liver development, p53 promotes a p21-induced cell cycle arrest in polyploid hepatocytes^29^. Intriguingly, *p21*^−/−^ mice showed a significant increase in ploidy, compared to WT animals. Curiously, the percentage of 4N nuclei in *p21*^−/−^ mice was still somewhat lower compared to that seen the *PIDDosome* knockout mice (∼16 % *vs.* ∼26%, respectively), suggesting that p21 and the PIDDosome may act in separate pathways (Figure 4D). However, comparing CM nuclei from *Casp2*-deficient mice with those from *Casp2/p21* double-knockout (dKO) animals did not reveal an additional ploidy increase that was statistically significant, suggesting that p21 acts downstream of the PIDDosome (Figure 4D). Since p73 has been implicated in regulating CM proliferation and can also be regulated by MDM2 in absence of p53^83,84^, we reasoned that it may replace p53 during postnatal heart development. Thus, we have analyzed the CM nuclear ploidy of *p73* knockout mice at P7, a time when the PIDDosome is already active. However, no difference in tetraploid CM population was evident between *p73^−/−^* and WT mice (Suppl. Figure 3C), indicating that p73 does not compensate for p53 during heart development, as seen in other developmental settings^92^. Please note that we were unable to obtain viable *p73^−/−^* pups on P14.

In an attempt to identify alternative substrates processed by Caspase-2 in response to PIDDosome activation in CMs, we performed N-terminomics hearts isolated from *XMLC2^-^ Casp2^fl/fl^* and *XMLC2^+^Casp2^fl/fl^*mice on P7 (Figure 4E and Suppl. Figure 3D). This analysis identified a total of 7503 unique neo N-termini, of which 547 peptides resulted from a cleavage after aspartate residues, a hallmark of caspase-mediated proteolytic activity (Table S1). Importantly, most of the cleavage events after aspartic residues were enriched in the control samples that express Caspase-2 (*XMLC2^-^Casp2^fl/fl^*) proving the validity of the approach. Among the putative Caspase-2 targets, two proteins were of interest: myosin heavy chain (MYH7), and Prothymosin alpha (PTMA). MHY7 is more abundant in fetal, compared to adult hearts^59,63^. In the absence of Caspase-2, processed MHY7 is less abundant, suggesting that, in line with the observed transcriptomic changes (Figure 4C), Caspase-2 loss delays CM maturation. Intriguingly, PTMA, the second substrate of interest, reportedly promotes CM proliferation^93^ and caspase-3-mediated proteolysis of PTMA has been noted in apoptotic cells^94^. Thus, the increased abundancy of unprocessed PTMA in Caspase-2-deficient CMs might facilitate prolonged proliferation and thus, an increased CM ploidy.

Taken together, our findings support a role for p21 downstream of the PIDDosome that regulates CM ploidy independent of p53. If Caspase-2-mediated proteolysis of MHY7 and/or PTMA are indeed critical for ploidy control downstream of the PIDDosome needs to be addressed in future studies.

## 4. Discussion

During early postnatal development, cardiomyocytes lose their proliferative capability and become polyploid, which is an established barrier to heart regeneration^11,12^. To date, little is known how CM polyploidy is established in vertebrates, which molecular pathways control this process during development and why polyploid cardiomyocytes cannot re-enter the cell cycle.

Here, we show that the PIDDosome multiprotein complex is critical for regulating cardiomyocyte polyploidization. In fact, mice lacking individual PIDDosome components exhibit an increased nuclear and cellular ploidy status in cardiomyocytes, which does not alter cardiac function nor structure, at least in steady-state (Figure 4F). If loss of PIDDosome function may affect tissue performance after damage e.g., after myocardial infarction, needs to be tested experimentally. Yet, the PIDDosome represents an attractive target to modulate cell cycle arrest in polyploid CM, and thus, we could speculate that its suppression might be beneficial for cardiac regeneration after heart injury. Consistent with timed DNA content increase during postnatal heart development^7^, the PIDDosome-mediated and cardiomyocyte-specific ploidy control is restricted to the first two weeks of life. Of note, the relative expression of the PIDDosome components also increase during that time and drop once CMs became polyploid^7^, a finding reminiscent of observations made in the liver^23^. Similarly, as in hepatocytes, restriction of polyploidy is dependent on the ability of PIDD1 to localize to extra centrosomes, supported by findings in ANKRD26-deficient hearts. Interestingly though, in CMs the PIDDosome limits polyploidy by modulating different cell cycle genes, including *p21,* but independently of p53 activity. Interestingly, CyclinG1, a p53 target gene, was shown to affect ploidy levels in the heart when overexpressed in neonatal rat CM, while its loss in mice prevented ploidy increases in response to pressure overload. However, while p53 has been repeatedly implicated in CM death in response to DNA damage, a direct impact on CM development has never been reported. How the PIDDosome eventually engages p21 remains unresolved. Yet, as *Casp2/p21* double-mutant animals show no additional ploidy increase over that seen in *Casp2*-deficient mice, we conclude that *p21* is epistatic downstream of Caspase-2. How Caspase-2 can engage p21 in the absence of p53 is unclear, but stabilizing effects of Caspase-2 on *p21* mRNA translation have been reported^85^. Regardless of mechanism, these data together with the increased percentage of 4N cardiomyocyte nuclei in *p21* knockout animals suggest that the PIDDosome might exert its function not exclusively via p21, as the increase in 4N cardiomyocyte nuclei in *p21* knockout mice was significantly lower than that seen in PIDDosome-deficient CMs. These findings are in line with previous studies which have shown that postnatal cardiomyocyte proliferation depends on p21^86^ but not on p53, as a *p53* knockout is insufficient to induce cardiomyocyte proliferation^87^.

E2F8 is known to be a fundamental and positive regulator of hepatocyte ploidy during liver development^80^ where it transcriptionally targets *Casp2* and *Pidd1*^23^. Recently, it was reported that cardiomyocyte-specific double knockout of *E2F8* and *E2F7* mice have an increased percentage of mononucleated diploid cardiomyocytes, which had no effect on cardiac function in steady state and also failed to improve heart regeneration after infarction^88^. Our bulk nuclear RNAseq data has shown that cardiac-specific deletion of Caspase-2 correlates with an upregulation of *E2f8,* which could explain the increased level of polyploid cardiomyocyte in these mice^88^. *E2f7/8* double knockout livers are characterized by almost exclusively mononucleated diploid hepatocytes, while in the heart the inactivation of *E2f7/8* causes a defined increase of mononucleated diploid cardiomyocytes from ∼1% to 5%, but not more^88^. Thus, comparing liver and heart development, it seems that there are additional mechanisms which restrict cardiomyocyte polyploidization during heart development aiming to secure steady state heart function. Finally, heart regeneration therapies based on cardiomyocyte proliferation face many obstacles in clinical studies and hence, dissecting the events that regulate polyploid cardiomyocyte proliferation during postnatal heart development will be essential for translational medicine. As such, the PIDDosome represents an attractive target to modulate cell cycle arrest in polyploid CMs. Of notice, small molecules targeting Caspase-2 have been reported^89–91^. Therefore, our findings not only describe a critical regulatory step in the terminal differentiation program activated postnatally in CMs but also open a new perspective for regenerative therapies based on polyploid CMs, instead of focusing on the limited regenerative potential of the few mononucleated CM proliferation present in the mammalian heart.

## Supporting information

N-terminomcs results

## Funding

ML acknowledges support by the Austrian Science Fund (FWF) (Lisa Meitner, M 3115-B), AV acknowledges support by the FWF (P36658, I6642), as well as the ERC, AdG 787171 (POLICE). FE acknowledges support from the DOC fellowship program of the Austrian Academy of Sciences (ÖAW). KM and MB acknowledges support by the CRC/TRR353 (Death Decisions, SP02) and CRC1453 (NephGen) funded by the German Research Foundation (DFG), and the German Federal Ministry of Education and Research (BMBF) within the Medical Informatics Funding Scheme, PM4Onco (FKZ 01ZZ2322A). The Orbitrap Eclipse mass spectrometer was funded in part by the German Research Foundation (INST 95/1650-1). JM also acknowledges support by the CRC/TRR353 (Death Decisions, SP01), funded by the German Research Foundation (DFG).

## Acknowledgements

We would like to thank Claudia Soratroi, Irene Gaggl and Julia Heppke for technical support, Martin Saurwein and Maria Fischer for animal care, as well as Bergmann Olaf, TU-Dresden, DE, for critical discussion and Andrew J Holland (Baltimore) for heart tissue from *Ankrd26^−/−^* mice, Brigitte Jenewein for supporting with imaging acquisition and Lexogen for support with nuclear RNAseq analyses. The images were created by BioRender, https://BioRender.com/t34z245

## Conflict of Interest

None declared.

---

**Supplemental Figure 1. CRE expression under control of the *XMLC2* promoter does not affect CM nuclear ploidy.**

(**A**) Schematic representation of the flow cytometry gating strategy used to assess CM nuclear ploidy. Nuclei were isolated from frozen hearts and CM nuclei were identified as PCM1^+^. The DNA content was measured by PI staining (**B-C**) Schematic illustrations of the mouse generation of cardiomyocyte-specific deletion of *Casp2* (B) and cardiomyocyte-specific membrane labelling (mT/mG)(C) driven by *XMLC2*-dependent Cre activation. (**D**) Quantification of CM nuclear ploidy of the indicated genotypes measured by flow cytometry. n: 3 (*XMLC2^-^*), 4 (*XMLC2^+^*). Data are mean ± SD analyzed by Two-way ANOVA; n.s.: not significant. (**E**) Representative H&E stained FFPE heart sections of 3 months-old mice of the indicated genotypes. Scale bars: 1mm.

**Supplemental Figure 2. Gene set enrichment analysis results of control CMs supports the previously described developmental changes between P1, P7 and P14 CMs.**

**(A**) RLDF (regularized linear discriminant functions) representation of the CM RNAseq samples, based on the centered, log2-transformed counts-per-million (log2CPM) (note: before count normalization with limma). The sample groups as indicated in the legend were defined as RLDF training groups. RDLF was performed with the function ‘plotRLDF’ of the R package limma (v. 3.52.4). (**B**-**C**) Selected significantly deregulated gene sets of the gene set enrichment analysis results comparing developmental days in control mice (*XMLC2^-^ Casp2^fl/fl^*). Dashed lines represent a plain p-value of 0.05. (**D**) Heatmap of selected genes involved in sarcomere structure and regulation. The Z-scores of normalized gene expression calculated across P1 and P7 control samples are shown.

**Supplemental Figure 3. Bulk RNA sequencing sample exclusion**

(**A**) PCA of the CM RNAseq samples, based on the centered, log2-transformed counts-per-million (log2CPM) (note: before count normalization with limma). The samples are labelled with their respective group and replicate number; for improved readability of the labels, ‘*XMLC2^-^ Casp2^fl/fl^*’ was replaced with ‘minus’ and ‘*XMLC^+^ Casp2^fl/fl^*’ was replaced with ‘plus’. (**B**) Top 20 enriched gene sets (by adjusted p-value) of fuzzy clusters 6 and 7 genes determined by functional enrichment analysis. (**C**) Quantification of the CM nuclear ploidy of *p73^−/−^* mice measured by flow cytometry at P7. n:4 (WT and *p73^−/−^*). Data are mean ± SD analyzed by Two-way ANOVA; n.s.: not significant. (**D**) Principal component analysis based on log2 transformed N-terminal peptide intensities.

**Supplemental Figure 4. Fuzzy clustering of all samples**

Fuzzy clustering of all CM RNAseq samples. Clustering was performed on the gene log2 fold changes with respect to the average gene counts of the four P1 *XMLC2^-^ Casp2^fl/fl^* replicates, after each sample’s raw counts were normalized with the average count of its interquartile range count values.

## Notes

### Competing Interest Statement

The authors have declared no competing interest.

### Summary of Updates

We have included additional data, such as N-terminomics, aiming to identify Caspase-2 substrates, we also analyzed p73 KO mice. As a result, we also included additional authors

