## Supplementary figures and images for "The PIDDosome controls cardiomyocyte polyploidization during postnatal heart development"

### Supplemental Figures

#### Leone M *et al.* Suppl Figure 1

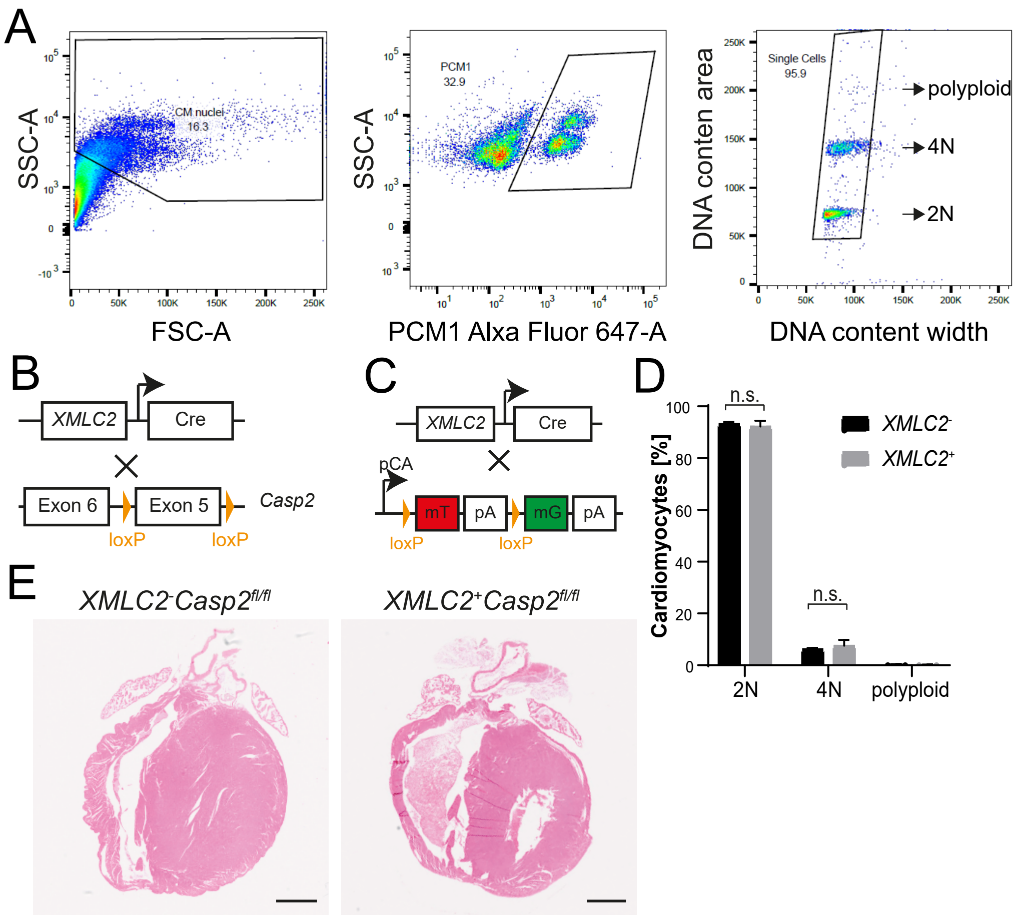

#### Leone M *et al.* Suppl Figure 2

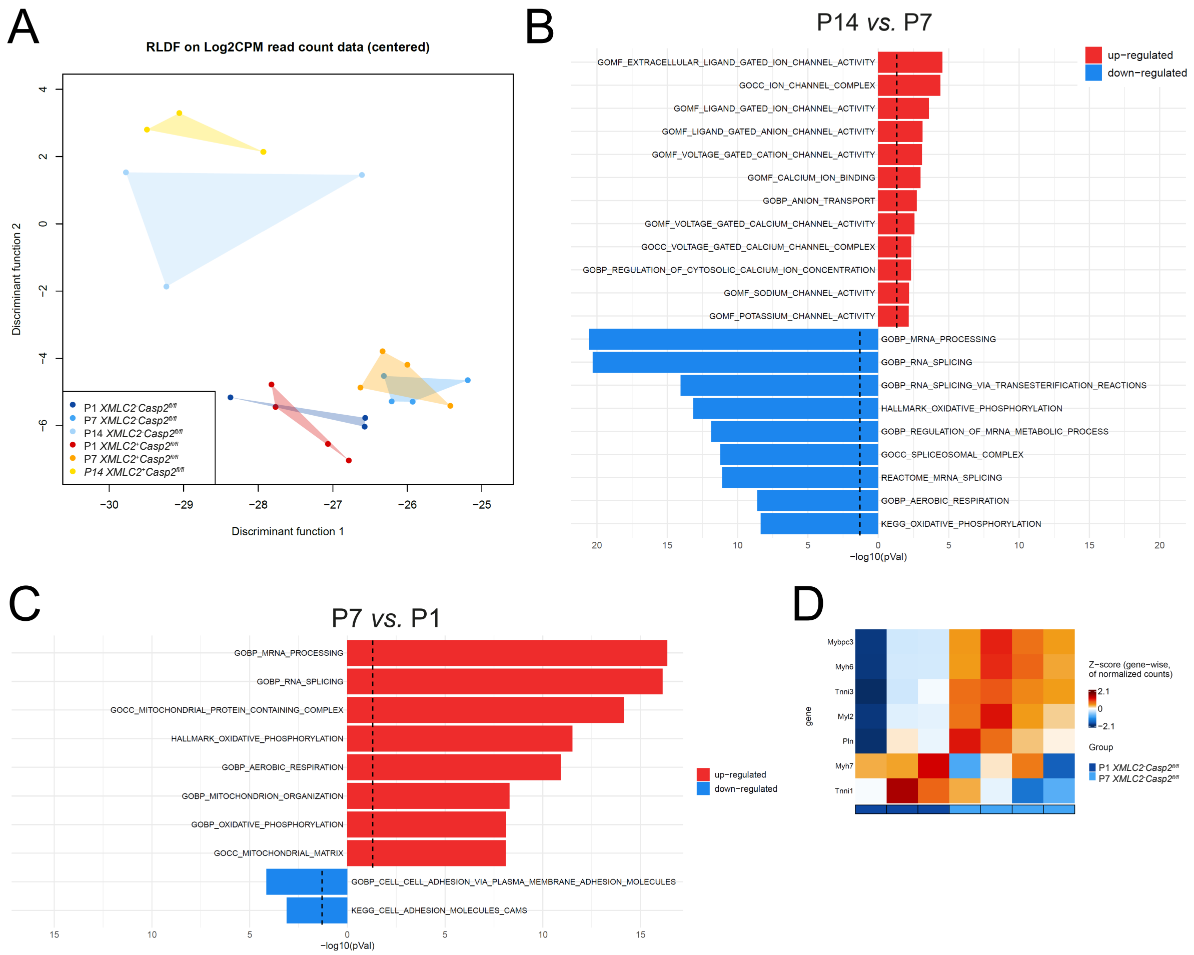

#### Leone M *et al.* Suppl Figure 3

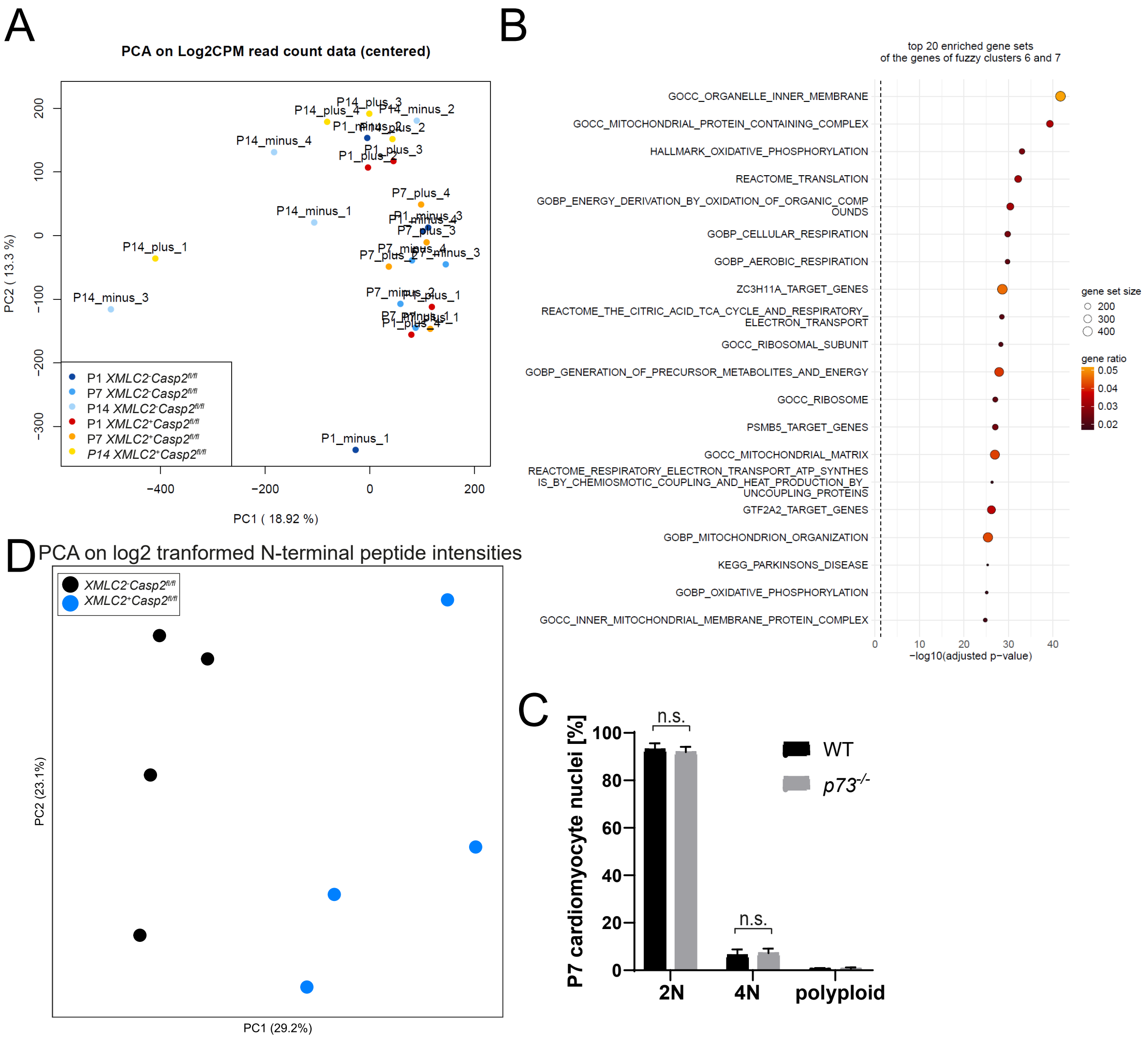

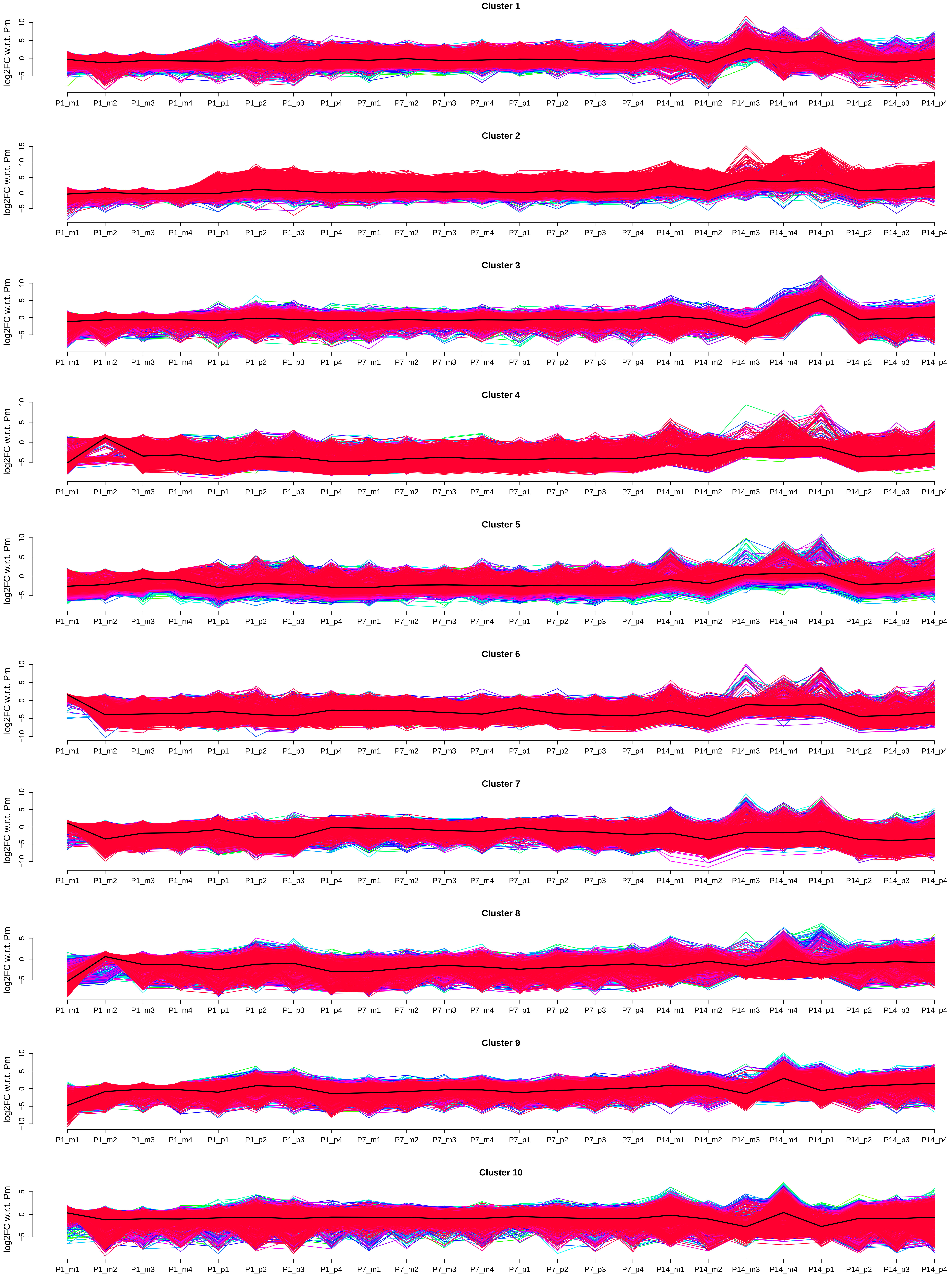
